# Patterns and Progress of Peninsular Malaysia’s Amphibian Research in the 21st Century (2000–2020)

**DOI:** 10.1101/2021.05.17.444585

**Authors:** Kin Onn Chan, Norhayati Ahmad

**Author notes:** Corresponding Author’s.

## Abstract

In this study, we review the status, patterns, and progress of Peninsular Malaysia’s amphibian research in the 21st century with the main goal of identifying areas for improvement that can help focus and prioritize future research initiatives. Between 2000–2020 we found 130 publications that can be broadly categorized into four groups: 1) checklists and biodiversity; 2) new species and taxonomy; 3) ecology and natural history; and 4) evolution and phylogenetics. An average of 6.5 papers was published per year and although the number of papers fluctuated, there was a significant upward trend in the number of papers published. Almost half (49%) of all papers published comprised checklists and biodiversity-related papers. This was followed by new species and taxonomy (25%, 33 papers), evolution and phylogenetics (14%, 18 papers), and ecology and natural history (12%, 16 papers). Amphibian research was conducted most frequently in the states of Kedah, Pahang, and Perak, and most infrequently in the states of Malacca, Negeri Sembilan, Selangor/Kuala Lumpur, Perlis, and Kelantan. Despite being a megadiverse country and a biodiversity hotspot, not a single conservation-centric paper has ever been published on Peninsular Malaysian amphibians, highlighting the urgent need for future research to focus on conservation.

## Introduction

Amphibian research in Peninsular Malaysia began in earnest towards the late 19^th^ and early 20^th^ century with notable contributions from early explorers (e.g. Ferdinand Stoliczka), herpetologists (e.g. Malcolm Arthur Smith and Norman Smedley), and workers at the British Museum (e.g. George Albert Boulenger). Understandably, early collections were centered around port and administrative cities in the states of Penang (Stoliczka 1870), Malacca (Boulenger 1887), and Perak (Boulenger 1900; Smith 1935), as well as important Hill Stations such as Cameron Highlands (Smedley 1931). Additionally, several expeditions to high elevation mountains were carried out (Boulenger 1908; Dring 1979; Grandison 1972). These early collections resulted in a throve of new species and laid the groundwork for a better understanding of Peninsular Malaysia’s amphibian fauna in the 21^st^ century.

At the turn of the 21^st^ century, important contributions continue to be made, largely led and inspired by the prominent Malaysian zoologist Dr. Lim Boo Liat (Das *et al.* 2004; Grismer *et al.* 2001, 2004; Kiew *et al.* 1995; Leong & Lim 2004; Lim *et al.* 2002; Norhayati *et al.* 2005). Alongside surveys, inventories, and new species discoveries that continue to be forthcoming, the breadth of research topics also expanded to include research in the field of molecular systematics and evolution (Chan *et al.* 2017, 2018b, 2020c, b; a; Chan & Brown 2017, 2020; Chan & Grismer 2019; Davis *et al.* 2016; Hurzaid *et al.* 2014; Matsui *et al.* 2017; Stuart *et al.* 2006). As research continues to develop with changing times and technologies, we review the status, patterns, and progress of Peninsular Malaysia’s amphibian research in the 21st century with the main goal of identifying areas for improvement that can help focus and prioritize future research initiatives.

## Material and Methods

We performed a comprehensive literature review of amphibian research publications in Peninsular Malaysia from the year 2000–2020. Only peer-reviewed primary literature was included and studies that did not directly involve material from Peninsular Malaysia were not considered. In total, 130 publications were found and categorized into the following groups:

1) Checklists and biodiversity: surveys, inventories, checklists, distribution, and measures of diversity.
2) New species and taxonomy: new species descriptions, taxonomic notes, and revisions.
3) Ecology and natural history: natural history notes, behavior, disease, and environmental correlates.
4) Evolution and phylogenetics: evolutionary history and phylogenetic relationships.

To characterize research trends and patterns, we analyzed the dataset according to the year of publication, number of papers, category, and geography. For geography, we classified papers according to states. Studies that were not focused on a particular state were excluded. All analyses were performed in R (R Core Team 2014). The categorized bibliography of all papers used in this study is presented in the Appendix.

## Results

Despite fluctuations from year to year, the 20-year trend showed a significant increase in the number of papers published (Fig. 1). On average, 6.5 papers were published per year, with a median of 5.5. The lowest number of papers published was one (years 2001 and 2005), while the highest number was 17 (year 2020).

**Fig. 1.**
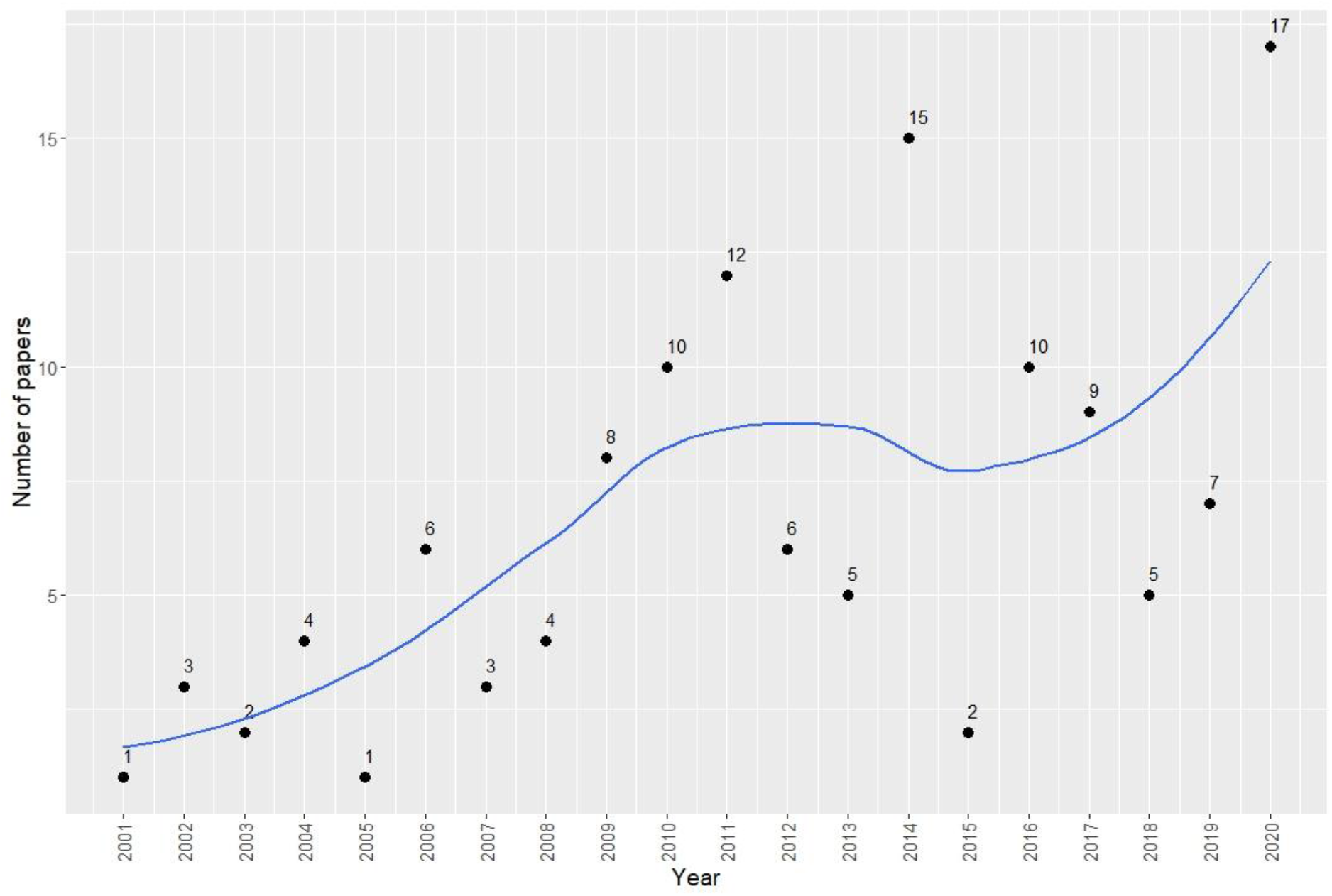
Scatter plot showing the number of amphibian research papers published by year. The blue line represents the regression line obtained using a local polynomial regression fitting method.

Almost half (49%) of all amphibian research published between 2000–2020 was represented by checklists and biodiversity-related papers (Fig. 2). This was followed by new species and taxonomy (25%, 33 papers), evolution and phylogenetics (14%, 18 papers), and ecology and natural history (12%, 16 papers). Checklists and biodiversity-related papers also dominated the research literature across all years (Fig. 3). New species and taxonomic papers ranked second highest from the years 2000–2015. Between 2016–2020, the number of ecology, natural history, evolution, and phylogenetic papers increased more than two-fold and outnumbered papers on new species and taxonomy (Fig. 3).

**Fig. 2.**
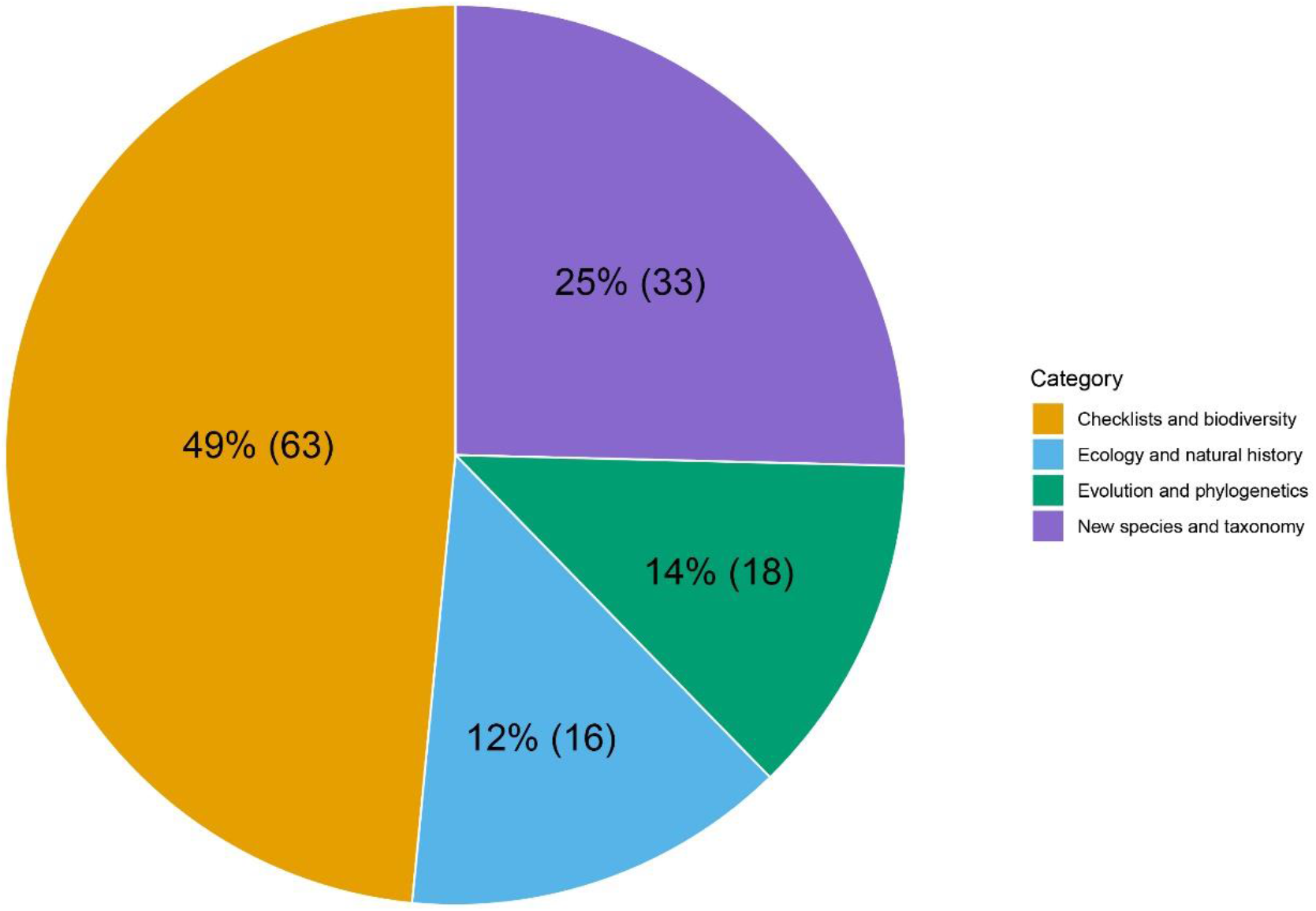
Percentage and number of papers published (in parenthesis) by category between 2000–2020.

**Fig. 3.**
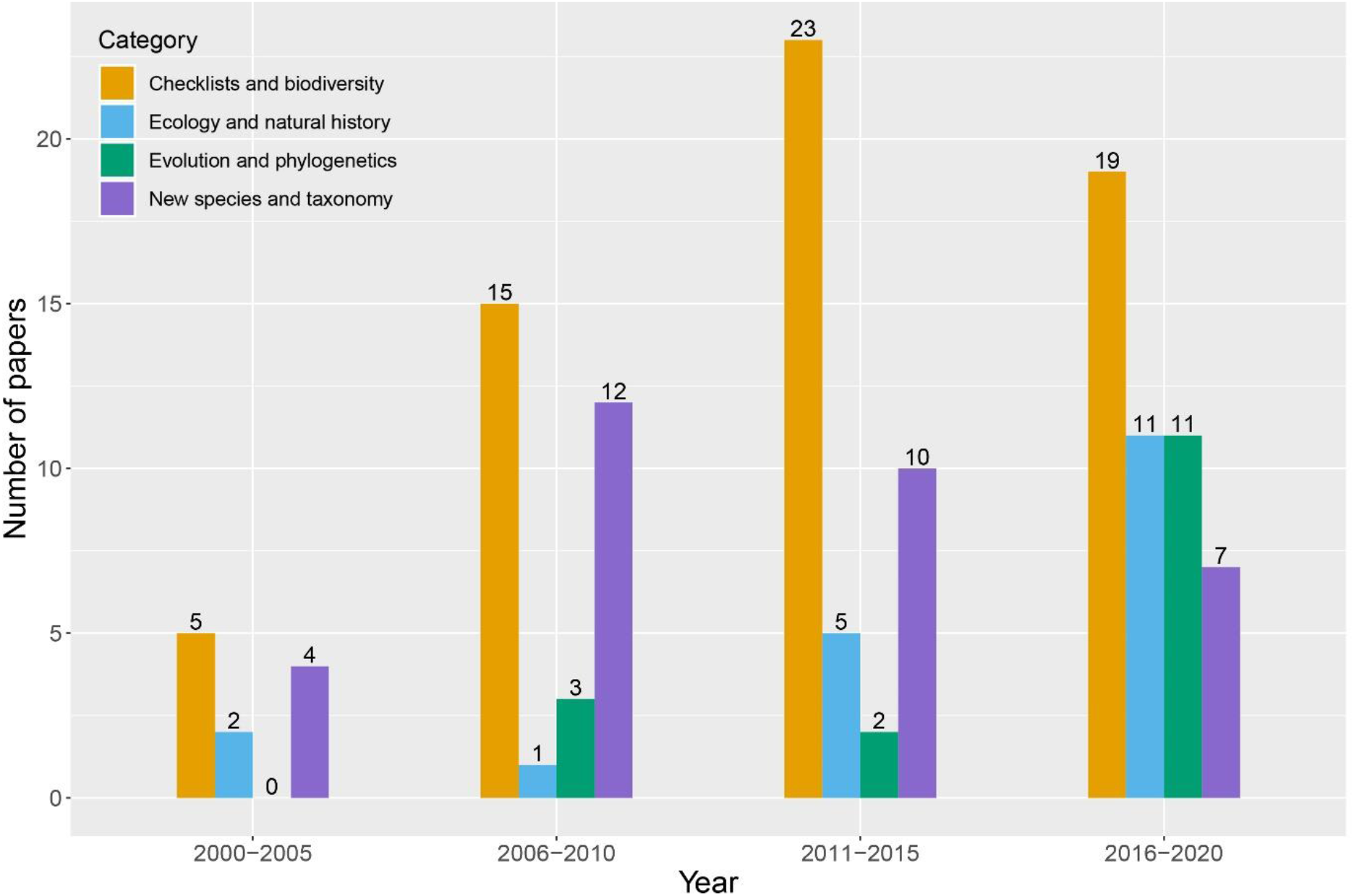
Bar plots grouped by category and binned into 5-year periods. Numbers above bars represent the total number of papers published in each category.

Amphibian research was conducted most frequently in the states of Kedah, Pahang, and Perak, and most infrequently in the states of Malacca, Negeri Sembilan, Selangor/Kuala Lumpur, Perlis, and Kelantan (Fig. 4).

**Fig. 4.**
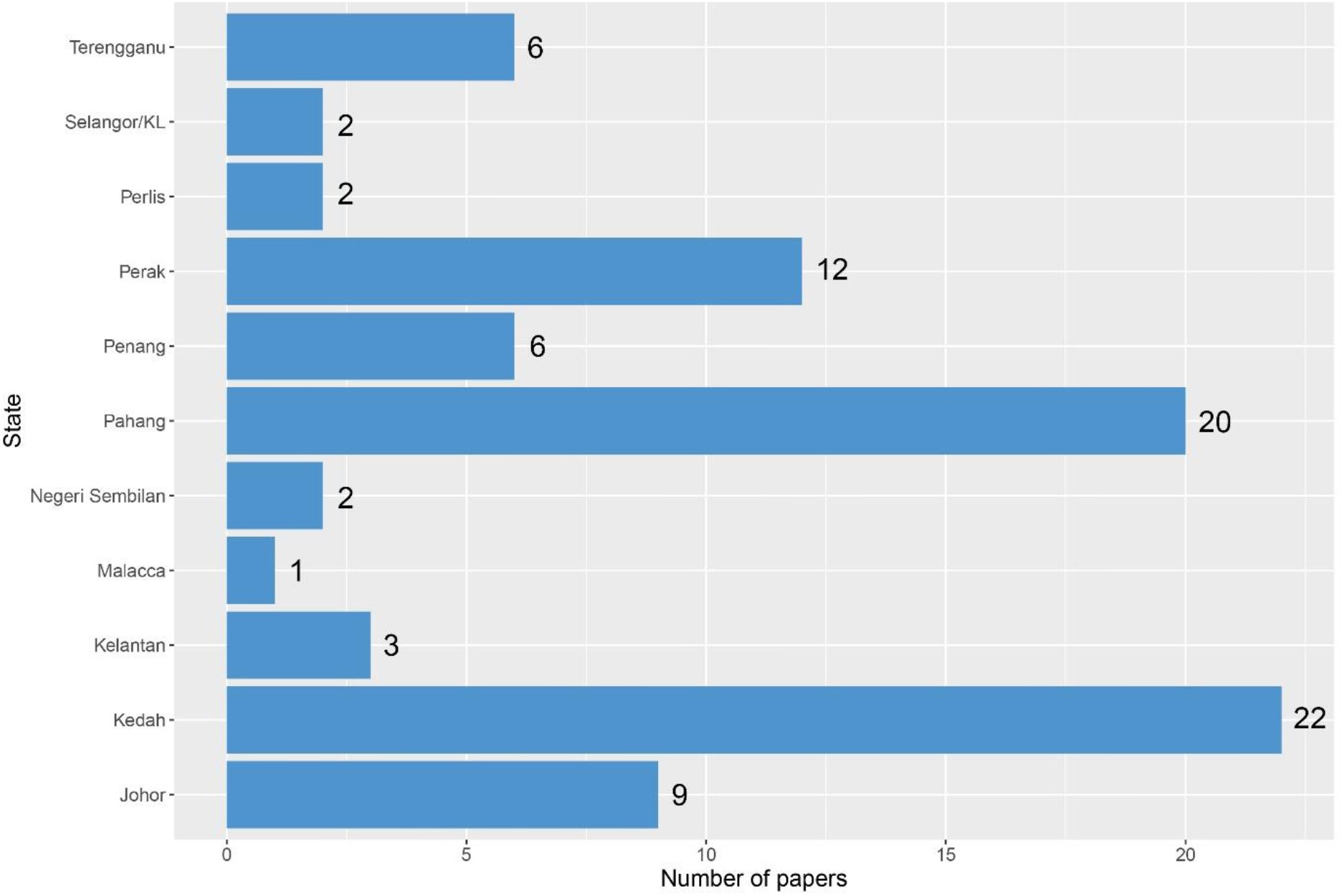
Number of papers published by state. Only studies that were explicitly conducted within a particular state were included.

## Discussion

The increasing number of papers published per year shows that amphibian research in Peninsular Malaysia is on an upward trend and the dominance of checklists, biodiversity, new species, and taxonomy-related papers indicate that the biodiversity of amphibians in Peninsular Malaysia is still far from being fully realized. The number of papers published by state (Fig. 4) shows that some regions are still poorly studied. This most notably includes the states of Perlis, Negeri Sembilan, Malacca, Kelantan, Selangor/Kuala Lumpur, and Terengganu. Research in some of these areas has resulted in numerous important discoveries (Chan *et al.* 2011, 2014b, 2018a), indicating that these areas are still poorly studied and should be prioritized for future research. However, states that have received more research attention such as Perak, Pahang, and Kedah continue to produce new discoveries (Chan *et al.* 2014a; Davis *et al.* 2016; Quah *et al.* 2011) and thus, research intensity in these states should continue and not be reduced.

Over the years, the number of papers on ecology, natural history, evolution, and phylogenetics has increased, particularly in the last five years (Fig. 3). In comparison, not a single paper on evolution and phylogenetics was published between 2000–2005. This could be due to the lack of expertise and the high cost of genetic sequencing during that period. Advances in sequencing technology coupled with a concomitant reduction in sequencing cost and collaborations with international partners saw a rise in genetic research between 2005– 2020. These, including more recent studies involving genome-scale data (Chan *et al.* 2020a, c; b) indicate that amphibian research in Peninsular Malaysia is progressing and is on par with other research programmes around the world.

One notable research gap is the lack of conservation-based research. Despite being a megadiverse country and a biodiversity hotspot, not a single conservation-centric paper has ever been published on Peninsular Malaysian amphibians. Research in this field is urgently needed as Malaysia is one of the countries with the highest rate of deforestation (Hansen *et al.* 2013) with almost 30% of its amphibian species threatened (MyBis 2021).

## Appendix

## Checklists and Biodiversity

## Ecology and Natural History

## New Species and Taxonomy

## Evolution and Phylogenetics

